# Lipid nanoparticle-mediated delivery of mRNA into the mouse and human retina and other ocular tissues

**DOI:** 10.1101/2023.07.13.548758

**Authors:** Cheri Z. Chambers, Gillian L. Soo, Abbi L. Engel, Birth Defects Research Laboratory (BDRL), Ian A. Glass, Andrea Frassetto, Paolo G. V. Martini, Timothy J. Cherry

## Abstract

**Purpose:** Lipid nanoparticles (LNPs) show promise in their ability to introduce mRNA to drive protein expression in specific cell types of the mammalian eye. Here, we examined the ability of mRNA encapsulated in lipid nanoparticles (LNPs) with two distinct formulations to drive gene expression in mouse and human retina and other ocular tissues.

**Methods:** We introduced mRNA carrying LNPs into two biological systems. Intravitreal injections were tested to deliver LNPs into the mouse eye. Human retinal pigment epithelium (RPE) and retinal explants were used to assess mRNA expression in human tissue. We analyzed specificity of expression using histology, immunofluorescence, and imaging.

**Results:** In mice, mRNAs encoding GFP and ciliary neurotrophic factor (CNTF) were specifically expressed by Müller glia and retinal pigment epithelium (RPE). Acute inflammatory changes measured by microglia distribution (Iba-1) or interleukin-6 (IL-6) expression were not observed 6 hours post-injection. Human RPE also expressed high levels of GFP. Human retinal explants expressed GFP in cells with apical and basal processes consistent with Müller glia and in perivascular cells consistent with macrophages.

**Conclusions:** We demonstrated the ability to reliably transfect subpopulations of retinal cells in mice eye tissues *in vivo* and in human ocular tissues. Of significance, intravitreal injections were sufficient to transfect the RPE in mice. To our knowledge we demonstrate delivery of mRNA using LNPs in human ocular tissues for the first time.

**Translational Relevance:** Ocular gene-replacement therapies using non-viral vector methods are of interest as alternatives to adeno-associated virus (AAV) vectors. Our studies show that mRNA LNP delivery can be used to transfect retinal cells in both mouse and human tissues without inducing significant inflammation. This promising methodology could be used to transfect retinal cell lines, tissue explants, mice, or potentially as gene-replacement therapy in a clinical setting in the future.

## Introduction

The retina plays an integral role in visual processing, and disorders of the retina are diverse in clinical presentation and etiology. Inherited retinal disorders (IRDs) are a heterogenous group of genetically inherited disorders that results in vision impairment or blindness. There are numerous types of IRDs, including retinitis pigmentosa (RP), choroideremia, Stargardt Disease, cone-rod dystrophy (CRD), and Leber Congenital Amaurosis (LCA). To date, over 270 genes have been identified and associated with retinal disorders^1^. Estimates of IRD prevalence vary but estimate that IRDs affect approximately 1 in 2000 individuals and over 5 million people worldwide, negatively impacting quality of life and causing significant economic burden^2–4^.

Despite notable advances in diagnostic capabilities with next-generation sequencing (NGS) and long-read sequencing (LRS), therapeutic options for IRDs remain limited. IRDs caused by monogenic, loss-of-function mutations are particularly attractive candidates for gene replacement therapies. Ocular immune privilege has made viral-mediated gene therapy strategies feasible^5,6^. Adeno-associated virus (AAV)-based gene augmentation therapies for IRDs have been most promising clinically, although lentivirus-mediated strategies have also been investigated^7,8^. In 2017, the FDA approved voretigene neparvovec, an AAV2-based therapy to treat RPE65-mediated LCA and the first gene-replacement therapy for an IRD^9^.

There are limitations with the current viral-vector mediated therapies. Transgenes larger than 4.7 kb are challenging to deliver efficiently in AAV systems^10^. There are also concerns that viral vectors can cause inappropriate and excessive immune system activation^11^. Additional considerations must be made for delivery of the genetic material via intravitreal versus subretinal injections, with intravitreal injections generally considered safer since this method does not directly damage or detach the retina. However, subretinal injections provide more direct access to the subretinal space, and therefore, the retinal pigment epithelium (RPE)^12^.

Lipid nanoparticles (LNPs) are an alternative, non-viral based method of transfecting retinal tissue. DNA delivery using LNPs has had limited success in non-dividing cells, such as retinal cells^13^. However, delivery of mRNA encapsulated in LNPs provides an alternative to AAV-based therapies that allows for delivery of larger transgenes in a system that has low immunogenicity. Two previous studies demonstrated the ability to deliver mRNA to the mouse retina with varying LNP formulations^14,15^. Subretinal injections of LNPs containing mRNA transfected mouse retinal Müller glia, RPE, optic nerve head, and trabecular meshwork, in an apolipoprotein adsorption and phagocytosis-independent manner. Notably, intravitreal injections in these studies did not lead to LNP penetration and transfection of the RPE^14,15^. Additionally, peptide guided LNPs have been used to deliver mRNA to photoreceptors, Müller glia, and RPE in rodents and non-human primate retinas^16^.

Due the unique combination of advancements in identifying genetic mutations that cause retinal degenerations such as IRDs, ocular immune privilege, and the use of mRNA therapeutics, it is an exciting time to explore mRNA LNP gene augmentation therapies for the retina. In this study, we investigated the ability of specifically formulated mRNA and LNPs to transfect mouse and human retinal cells. The mRNA sequences were optimized with chemically modified uridine to limit activation of the innate immune response, as well as to maximize protein expression. LNPs were designed to bind to receptors that induce endocytosis of the mRNA; mRNA LNPs that are not released from endosomes fuse with lysosomes for degradation. One of the challenges about using LNP to deliver genomic material is the adverse immune response; the LNP formulations used in this study were shown to be cleared from the liver and spleen significantly faster and more effectively than older, legacy LNP formulations^17,18^.

We aimed to characterize the ability of these engineered mRNA LNPs to transfect retinal cells in both *in vivo* and *ex vivo* systems. Intravitreal injections of enhanced GFP (EGFP) mRNA LNPs into the mouse vitreous chamber resulted in EGFP expression in the Müller glia and retinal pigment epithelium (RPE) of the retina. Additionally, EGFP mRNA was expressed in human retinal and RPE cells. To our knowledge, this was the first study that demonstrated *in vivo* transfection of mouse RPE using intravitreal injections, which have the advantage of being less invasive and safer than subretinal injections. Furthermore, we examined markers of acute inflammation post-delivery of mRNA LNPs, showed delivery of a therapeutic agent ciliary neurotrophic factor (CNTF), and performed *ex vivo* transfections of human retina and RPE. The results from this study advance our understanding of potential applications for mRNA LNP technology in the rodent and human retina.

## Methods

### Intravitreal Injections

All protocols used in this study received approval prior to the start of experiments by the Institutional Animal Care and Use Committee of Seattle Children’s Research Institute (Protocol 00050). Six-to seven-week-old male CD-1 mice (Charles River Laboratories, Wilmington, MA) were injected intravitreally with mRNA lipid nanoparticles (LNPs). Mice were anesthetized in an isoflurane chamber. A sterile, 30G needle (BD Biosciences, Franklin Lakes, NJ) was inserted at an approximately 45° angle along the limbus to create an initial puncture at the border of the cornea and sclera. A beveled, glass micropipette with a 5 μM tip (Clunbury Scientific, Bloomfield Hills, MI) was filled with either 1X PBS, pH 7.4 or mRNA LNPs, with 0.1% Fast Green FCF dye (Sigma-Aldrich, St. Louis, MO). A FemtoJet 4i electronic microinjection machine (Eppendorf, Hamburg, Germany) was used to deliver the reagents into the same initial puncture hole with the following parameters: injection pressure 290 hPa and compensation pressure 10 hPa. A maximum of 1 μL volume of mRNA LNP (1.8 mg/mL) was introduced into the vitreous cavity per injection. The amino lipid formulations used in these studies, 2T(Heptadecan-9-yl 8-((2-hydroxyethyl)(8-(nonyloxy)-8-oxooctyl)amino)octanoate) and 6T (2- (dinonylamino)-1-(4-(N-(2-(dinonylamino)ethyl)-N-nonylglycyl)piperazin-1-yl)ethan-1-one), were previously tested in rat, mouse, and non-human primate *in vivo* models^17,18^.

### Histology of Mouse Retinas

Mouse eyes were enucleated, small holes were created in the corneas, and then fixed in 4% paraformaldehyde (PFA) in 1X PBS, pH 7.4 for 1 hour at room temperature while gently shaking. Eyes were rinsed in 1X PBS, then dissection was performed to remove any extraocular tissue, most of the cornea, and the lens. Eyecups were then processed and equilibrated in increasing concentrations of 5, 15, and 30% sucrose in 1X PBS and then in a well-mixed 1:1 solution of 30% sucrose/optimal cutting temperature compound (OCT) (Sakura Finetek USA, Torrance, CA) before freezing rapidly on dry ice. Frozen eyecups were sectioned to 40 μM thickness at -20°C on a Leica CM3050 S Cryostat (Leica, Wetzlar, Germany) onto TruBond 380 adhesive slides (Electron Microscopy Sciences, Hatfield, PA), allowed to dry at room temperature for 1 hour, then frozen with desiccant at -80°C.

### Immunofluorescence of Mouse Retinas

Slides were washed once with 1X PBS, then incubated with blocking buffer (0.1% Triton X-100, 0.02% sodium dodecyl sulfate, and 1% bovine serum albumin in 1X PBS) for 30 minutes. Slides were incubated with DAPI (Invitrogen Cat# D1306) and primary antibodies for 4 hours: chicken anti-GFP (1:750, Thermo Fisher Scientific Cat# A10262, RRID:AB_2534023), rabbit anti-SOX9 (1:500, Millipore Cat# AB5535, RRID:AB_2239761), rabbit anti-RPE65 (1:500, Abcam Cat# ab231782), rabbit anti-CNTF (1:500, Proteintech Cat# 27342-1-AP, RRID:AB_2880848), rabbit anti-Iba1 (1:500, FUJIFILM Wako Shibayagi Cat# 019-19741, RRID:AB_839504), rabbit anti-interleukin-6 (1:500, Abcam Cat# ab7737, RRID:AB_306031), and Isolectin GS-IB_4_ from *Griffonia simplicifolia*, Alexa Fluor™ 647 Conjugate (1:500, Invitrogen Cat# I32450). Slides were washed with 1X PBS three times and incubated in fluorescently coupled Alexa Fluor secondary antibodies (Invitrogen) for 2 hours. Slides were washed three times in 1X PBS and mounted with Prolong Gold Antifade Mountant (Invitrogen #P36931).

### LNP Treatment of Cultured Human Retinal Pigment Epithelium (RPE)

Developing human eye tissue was received from the University of Washington Birth Defects Research Laboratory (BDRL) with ethics board approval and maternal written consent was acquired before specimen collection in compliance with federal and state regulations. RPE from 115 day developing eyes was isolated according to established protocol^19^ and cultured in 5% v/v FBS in RPE MEM-alpha media with Rock inhibitor (10 μM) for 1 week and switched to 1% v/v FBS RPE media^20^. The cells were cultured to confluence and passaged 4 times before plating to 8-chamber slides (Thermo Fisher Scientific Cat#177445) coated with Matrigel (Thermo Fisher Scientific Cat# CB-40230 or Corning Cat# 356230) at a cell density of 200,000 cells/well.

These were grown for 5 weeks before starting the treatment. EGFP mRNA in 2TLNP was added at a ratio of either 10 μL to 190 μL 1% RPE culture media (0.1 mg/mL), 25 μL LNP to 175 μL media (0.25 mg/mL), or 50 μL LNP to 150 μL media (0.5 mg/mL). For controls, we used media only or 10 μL diluent or 50 μL diluent to media in the same ratio as LNP. Each control and LNP treatment were done in duplicate. The treatment was left on cells for 2 hours at 37°C and then removed and replaced with media only overnight at 37°C. The cells were then washed with 1X DPBS, fixed in 4% PFA with 0.01% Triton-X for 10 minutes, switched to 4% PFA with sucrose for 5-10 minutes, then washed twice with 1X DPBS before immunofluorescence staining.

### LNP Treatment of Human Fetal Eye Cups

As above, developing human eye was received from BDRL and within 8 hours of collection, we dissected off the cornea and lens, with slight lateral cuts to open the eye cup slightly. Each eye cup was incubated in 100 μL DMEM/F12 (Gibco Cat#11330-032) + 10% FBS + 1% Pen/strep with 100 μL EGFP mRNA in 2TLNP (1:1 DMEM/F12 to LNP); this mixture was pipetted under the vitreous along the lateral slit as close to optic nerve as possible. Eye cups were incubated for 4 hours at 37°C. Then, the EGFP mRNA LNP and DMEM/F12 media mixture was removed, eye cups were washed gently with 500 μL 1X DPBS and put in 500 μL 4% PFA to fix over night at 4C. Then changed to PBS wash, and put in 5% sucrose for about 8hrs, then removed and added 15% sucrose overnight at 4C, then 30% sucrose for about 24h, and then put into OCT. Sections were made at 20-40um thickness for IHC staining.

### Immunofluorescence of Cultured Human RPE and Fetal Eyes

The fixed cells and eyecups were incubated in blocking buffer (0.1% Triton X-100 with 0.02% SDS and 1% BSA in 1X PBS) for 1 hour at room temperature with mild shaking. Primary antibodies used were the following: anti-GFP (1:100, ThermoFisher Cat# A10262, RRID:AB_2534023) with anti-ZO-1 (1:100, Abcam Cat#ab221547, RRID:AB_2892660), and DAPI (1:500 of a 1:1000 solution, Invitrogen Cat# D1302). The cells and eyecups were incubated in primary antibody solution with gentle shaking at 4°C, and then removed the next day with 3 washes of 1X PBS. Cells were incubated in secondary antibodies for 1 hour, which consisted of anti-chicken IgY (1:500, ThermoFisher Cat#A11039, RRID:AB_2534096) and anti-rabbit IgG (1:500, ThermoFisher Cat# A32795, RRID:AB_2762835).

### LNP Treatment, Histology, and Immunofluorescence of Adult Human Globes

Post-mortem adult human globes from a de-identified donor were obtained from Lions VisionGift (Portland, OR). The tissue was provided with the donor’s de-identified medical records including the following information: time and cause of death, post-mortem interval prior to cryopreservation of tissue, age, perceived race, and sex. The donor did not have a past medical history of ophthalmological conditions or interventions. Two globes from one donor were used for these experiments. Tissue biopsy punches (4 mm) were used to collect cross-sections of retina, RPE, and sclera. Tissue explants were taken of the fovea/macula and at the optic nerve head, as well as approximately 15 additional punches of the peripheral retina. Explanted retina/RPE punches were incubated for 4 hours at 37°C in DMEM with either diluent, EGFP mRNA encapsulated in 2TLNP, or EGFP mRNA encapsulated in 6T LNP to a final concentration of 0.1 mg/mL. Then, the DMEM/LNP mixture was removed and replaced with fresh DMEM for an additional 20 hours.

Tissue explants were washed 3 times with 1X PBS, then fixed with 4% PFA in 1X PBS for 1 hour at room temperature while gently shaking. Explants were placed in blocking solution (0.1% Triton X-100, 0.02% sodium dodecyl sulfate, and 1% bovine serum albumin in 1X PBS) for 1 hour. Retinal explants were incubated with DAPI (Invitrogen Cat# D1306) and primary antibodies for 3 days: chicken anti-GFP (1:750, Thermo Fisher Scientific Cat# A10262, RRID:AB_2534023), and rabbit anti-GFAP (1:500, Abcam Cat# ab7260, RRID:AB_305808). Slides were washed with 1X PBS three times and incubated in fluorescently coupled Alexa Fluor secondary antibodies (Invitrogen) for 3 hours. Tissue punches were washed three times in 1X PBS and mounted with Prolong Gold Antifade Mountant (Invitrogen #P36931).

### Therapeutic Targets Review

We utilized RetNet to identify genes and loci implicated in IRDs^1^. For our literature review of potential mRNA LNP therapeutic targets, we then examined gene expression in the peripheral retina from Table S2 of a published single-cell human retina and organoid database ^21^. This atlas lists the level of gene expression (normalized to 10,000 transcript counts per cell type) in various retinal cell types. We cross referenced the two databases to rank disease genes by level of expression in Müller cells (MC1, MC2, and MC3) and retinal pigment epithelium (RPE). Genes included are above gene expression level of 1. Supplemental information about diseases and affected proteins was drawn from the RetNet database. Additional information regarding gene reference and protein coding region size was drawn from the Ensembl genome browser^22^. Candidate targets were filtered to remove artifacts of rod RNA contamination by removing those candidates whose expression was 2x higher in rods than Müller glia or RPE.

### Microscopy, Image Analysis, and Quantification

Images of retinal cryosections were obtained using sequential scanning protocols on a Leica TCS SP5 confocal microscope. Maximum intensity projections (MIPs) and scale bars were created using ImageJ software (NIH, Bethesda, MD). Quantification of EGFP expression in RPE cells was performed using ImageJ software. Single-channel images of ZO-1 (tight junction marker used to denote edges of RPE cells in these analyses) and EGFP were converted to gray scale first and then threshold adjusted to match signal intensity. The threshold image was converted to binary, and separate ZO-1 and EGFP masks were created. Then, the ZO-1 binary mask was subtracted from the EGFP mask. The regions of interest (ROIs) were measured by adjusting the range of particle size to capture EGFP. Areas of no signal were used for generating the background with the manual draw tool and measured for area, min/max, and integrated density, similarly to the EGFP positive regions. Images and quantifications were saved for statistical analysis.

### Statistical Analysis

Statistical tests were performed using GraphPad Prism version 9 (GraphPad Software, San Diego, CA). A one-way ANOVA test with Tukey correction for multiple comparisons was done to compare EGFP expression in RPE cells treated with diluent or EGFP mRNA LNPs.

## Results

### Transfection of CD-1 Mouse Eyes with mRNA LNPs and Analysis of Cellular Expression

To assess the ability of mRNA encapsulated in LNPs to transfect retinal cells in an *in vivo* system, mRNA LNPs were delivered to CD-1 mice, chosen for the similarity of mouse retinal layers to the human retina. The basic experimental set-up was the following: intravitreal injection of mRNA LNPs were performed (one per eye) on anesthetized adult, CD-1 mice, then allowed to incubate for various time points between 48 hours and 2 weeks before harvest and subsequent analysis using immunofluorescence and confocal imaging (Figure 1). First, to determine the ability to successfully deliver mRNA LNPs to the mouse eye, EGFP mRNA encapsulated in LNPs of two lipid formulations (2T and 6T) was injected intravitreally. Cellular uptake of the 2T formulation is low-density lipoprotein receptor (LDLR) mediated, while the uptake of 6T is LDLR-independent. Eyes were harvested at various time points to assess duration of the EGFP expression. Confocal microscopy images of retinal cross-sections were analyzed for EGFP expression. EGFP is expressed in multiple cell types in the retina 48 hours (Figure 2A), 72 hours (Figure 2B), 96 hours (Figure 2C), 1 week (Figure 2D), and 2 weeks (Figure 2E) post-injection with EGP mRNA for both 2T and 6T lipid formulations. Additionally, outside of the retina, there was EGFP expression in the cornea, iris, and iridocorneal angle 48 hours post-injection with EGFP mRNA LNP, 2T formulation (Supplementary Figure 1).

**Figure 1.**
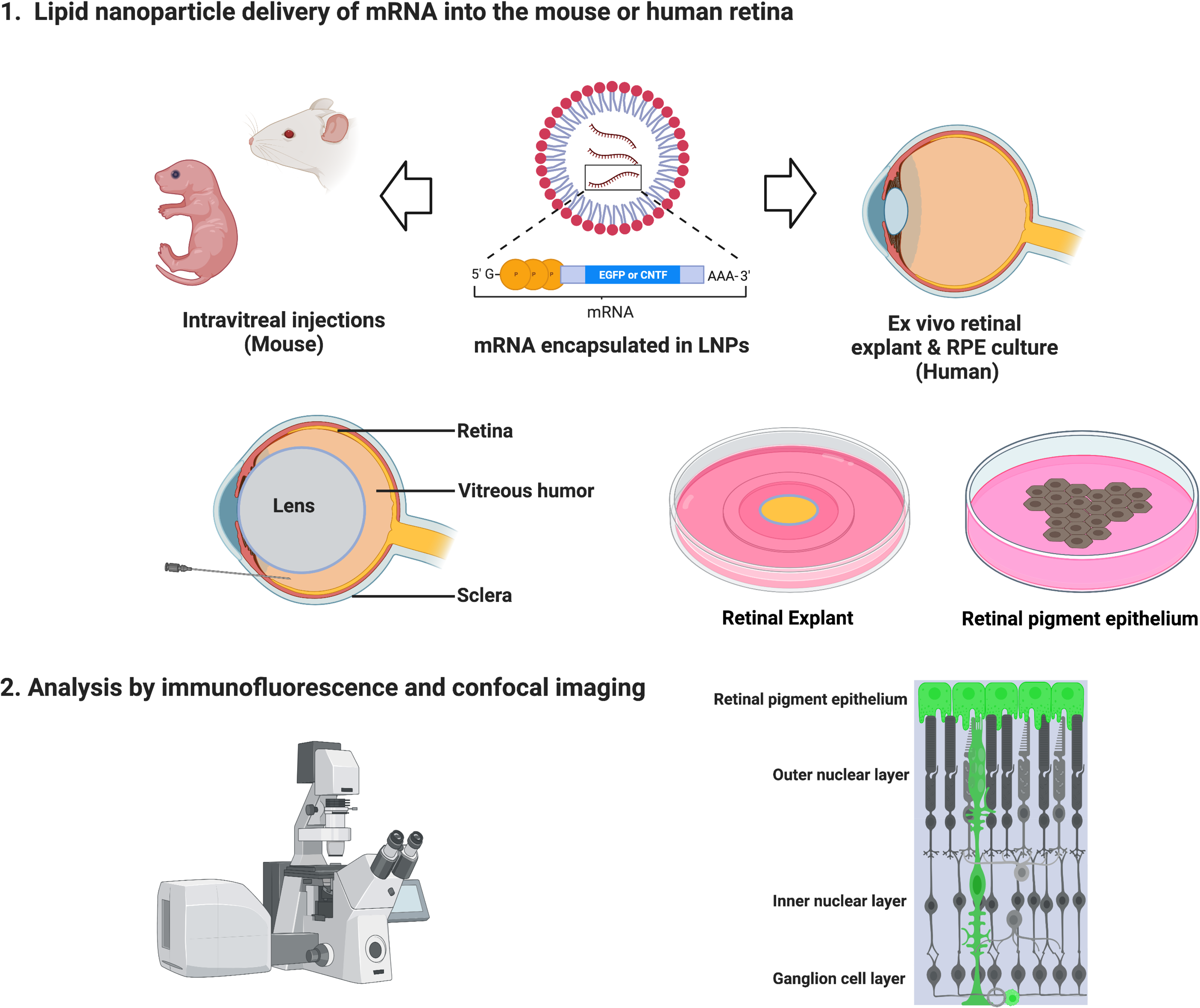
Schematic describes experimental workflow to deliver mRNA encapsulated in LNPs. CD-1 mice were used for *in vivo* experiments. Intravitreal injections of mRNA LNPs were performed to deliver the reagents to the vitreous chamber. After histological processing, analysis was completed using immunofluorescence and confocal imaging to determine transfection cell type expression and duration. Tile scan shows DAPI and GFP expression in a mouse retina cross-section 96 hours after intravitreal injection with EGFP mRNA encapsulated in 2TLNP. Representative image shown is a maximum intensity projection (MIP), scale bar: 500 μM.

**Figure 2.**
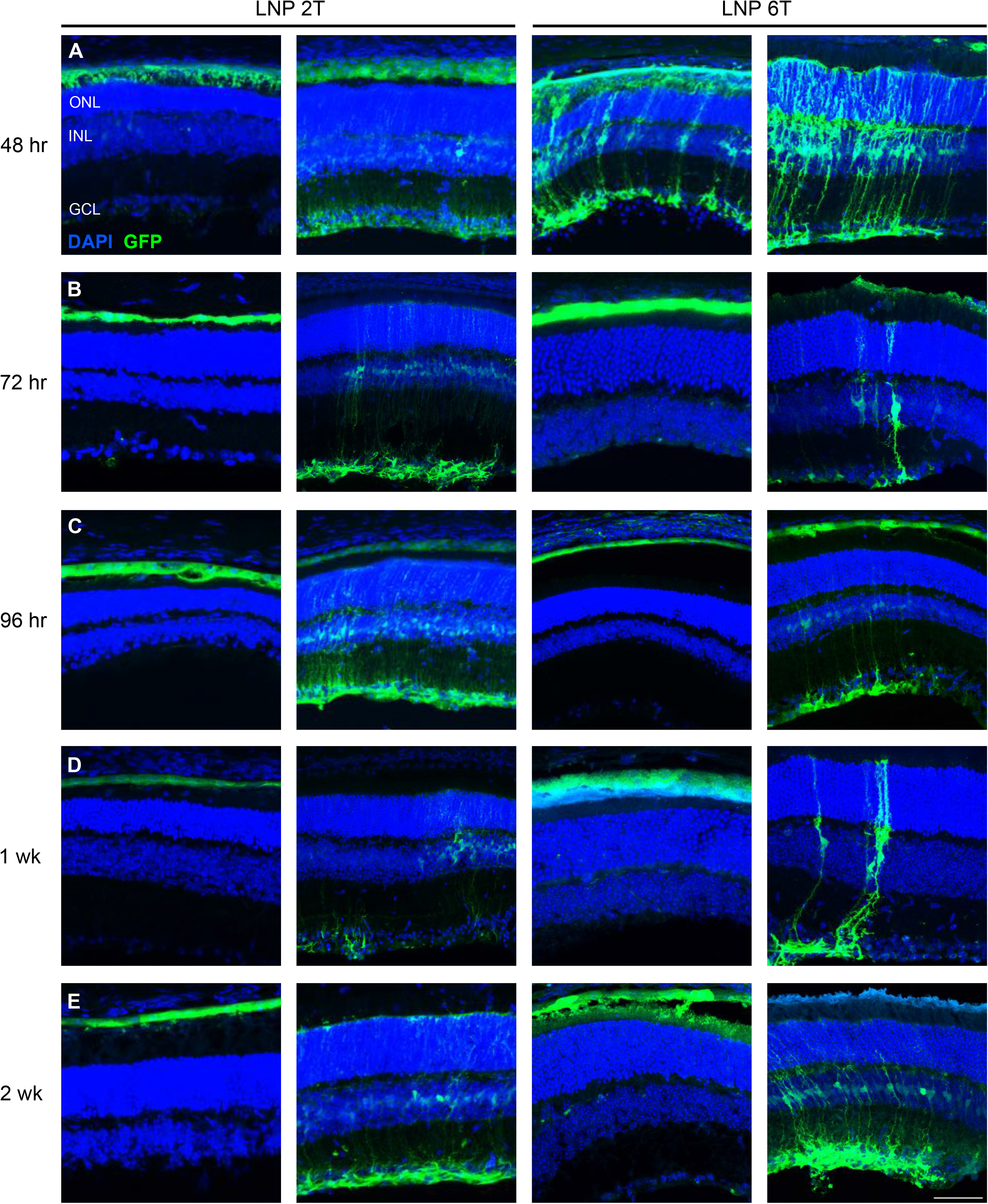
Intravitreal injections of EGFP mRNA transfects retinal cells in the mouse. Two amino lipid formulations, 2T and 6T, were compared for their ability to transfect retinal cells at various time points. Confocal images of retinal cross-sections with DAPI and EGFP staining are shown. EGFP is expressed in similar cell populations at (A) 48 hours, (B) 72 hours, (C) 96 hours, (D) 1 week, and (E) 2 weeks for both 2Tand 6T LNPs. Images shown are representative of at least 3 retinas and are MIPs (scale bar: 50 μM).

Next, to identify the specific retinal cell types transfected by EGFP mRNA LNPs, immunofluorescence using GFP and co-staining with antibody markers of Müller glia (anti-SOX9) and retinal pigment epithelium (anti-RPE65) was used in visualizing retinal cross-sections. Nuclei of Müller cells are stained with SOX9, which overlapped with nuclei transfected with EGFP for both 2T and 6T 48 hours post-injection (Figures 3A and 3C). Similarly, RPE cells are stained with RPE65, which overlapped with EGFP in the RPE of the retina for both lipid formulations (Figures 3B and 3D). Thus, EGFP was expressed in Müller glia and RPE cells in the mouse retina. Interestingly, the transfection pattern in neonatal CD-1 mice was partially different than in adult mice. EGFP mRNA 2TLNPs were injected intravitreally to P0 CD-1 pups, and retinas were harvested 24 hours after injection. Retinal cross-sections show EGFP expression in perivascular cells near endothelial cells (co-stained with anti-Isolectin GS-B_4_) and RPE cells (Supplementary Figure 2).

**Figure 3.**
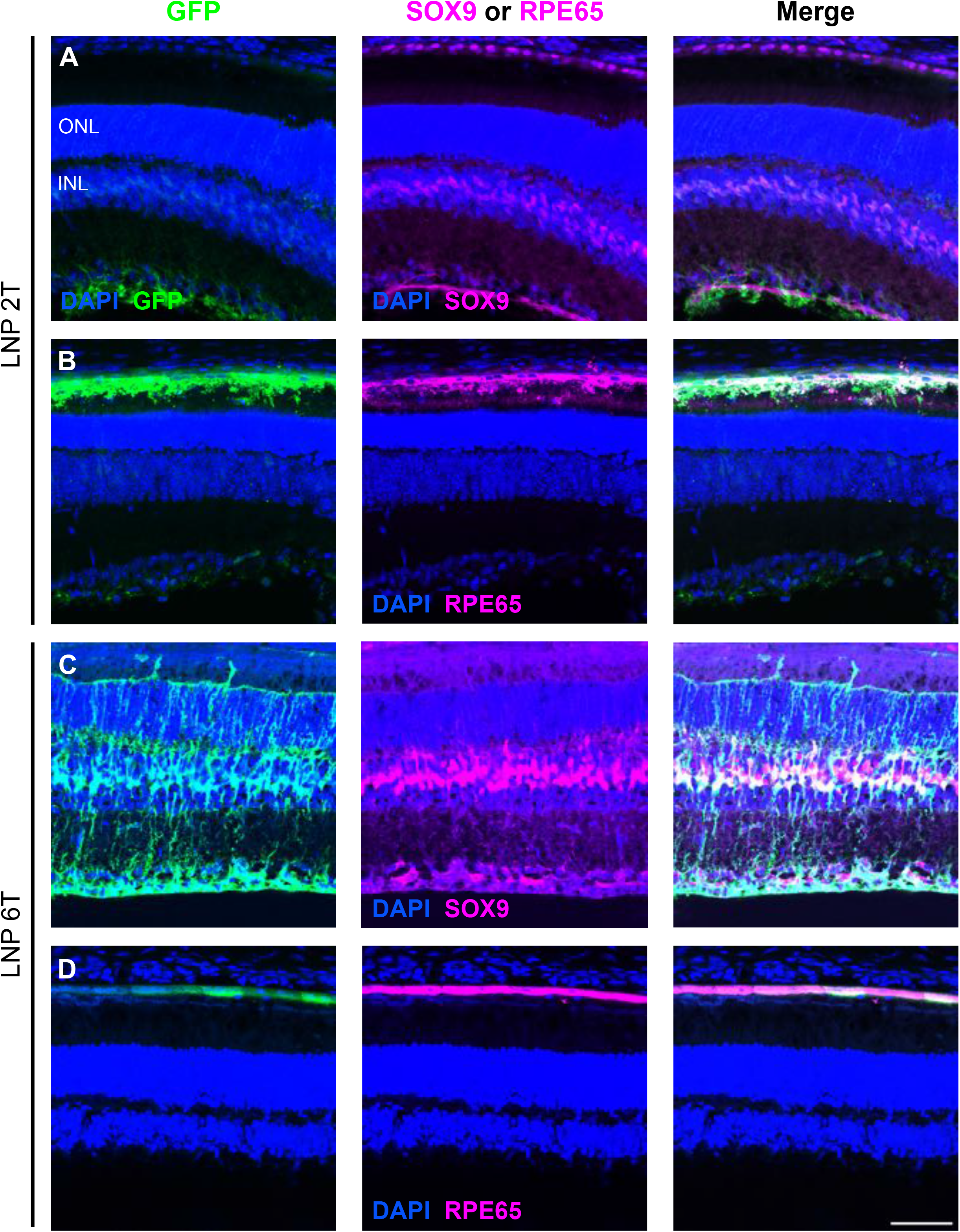
EGFP mRNA transfects Müller glia and RPE cells in the mouse retina. Retinal cross-sections were co-stained with DAPI, GFP, and either SOX9 or RPE65 48 hours after intravitreal injections of EGFP mRNA encapsulated in 2Tor 6T LNPs. (A) For the 2Tformulation, nuclei labelled with EGFP overlap with nuclei labelled with SOX9, indicating EGFP expression in the Müller glia. (B) Cells that express GFP also express RPE65, indicating EGFP expression in the RPE. Similarly, for 6T, there is co-expression of EGFP and (C) SOX9 and (D) RPE65. Images shown are representative of at least 3 retinas and are MIPs (scale bar: 50 μM).

Since the Müller glia and RPE cells can be transfected with EGFP mRNA LNPs, delivery of a potential retinal therapeutic agent, CNTF, was assessed next. EGFP mRNA 2TLNPs were delivered either alone via intravitreal injection or in conjunction with CNTF mRNA 2TLNPs in adult CD-1 mice. When EGFP mRNA is delivered alone, there was EGFP transfection of Müller glia, and CNTF was diffusely present in the retina, with slightly higher expression in the ganglion cell layer (GCL) (Figure 4A). Injection of EGFP and CNTF mRNA resulted in expression of EGFP in the Müller glia and increased expression of CNTF in the Müller glia that was absent in the EGFP mRNA alone condition (Figure 4B). This indicates that CNTF mRNA was successfully transfected into mouse retinal cells.

**Figure 4.**
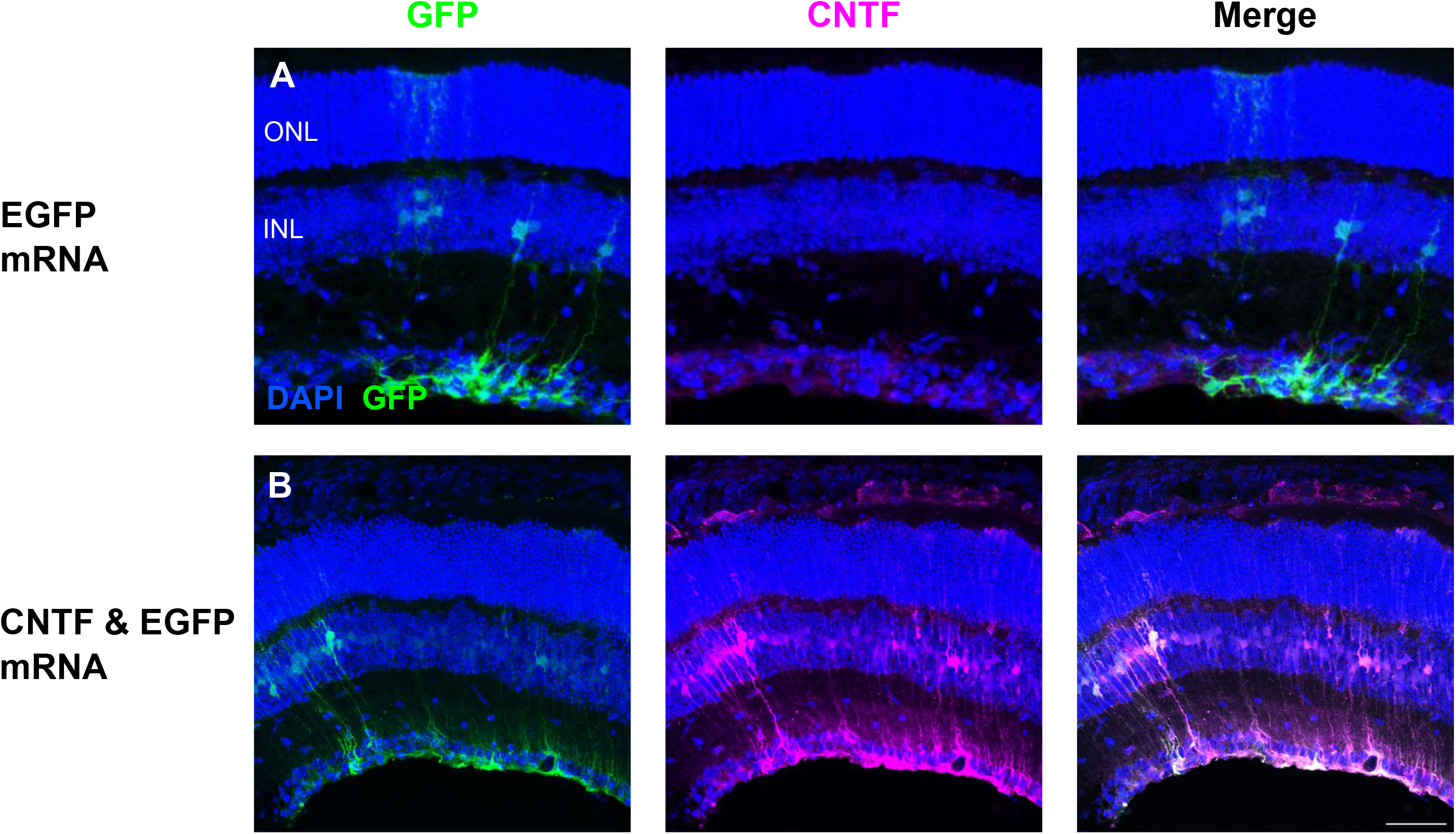
CNTF mRNA in 2TLNP transfects mouse Müller glia cells. Intravitreal injections of EGFP or both EGFP and CNTF mRNA were done. (A) Control EGFP injections show EGFP expression in the Müller glia and basal CNTF expression in the GCL based on immunofluorescence staining. (B) When EGFP and CNTF mRNA were co-delivered, EGFP mRNA expression is in the Müller glia. CNTF expression is increased and in Müller glia cells as well. There is only partial overlap of EGFP and CNTF expression, indicating that there are populations of Müller glia cells that express either EGFP only, CNTF only, or both. Images shown are representative of at least 3 retinas and are MIPs (scale bar: 50 μM).

### The Effect of mRNA LNP Delivery on Inflammation in the Mouse Retina

*In vi*vo inflammation in response to intravitreal injections of mRNA LNPs was investigated in adult CD-1 mice. Observation of mice post-injection yielded no external signs of inflammation (i.e., no conjunctival redness or discharge). A more detailed analysis was performed to compare levels and retinal locations of microglia (Iba1) and cytokine interleukin-6 (IL-6). Four conditions were compared for markers of inflammation: 1) no injection, 2) injection with 1X PBS, 3) injection with EGFP mRNA 2T LNP, or 4) injection with EGFP mRNA 6T LNP. Eyes were harvested 6 hours post-injection, and the distribution of microglia and IL-6 in the retinal layers was similar between all four conditions, indicating a baseline level of inflammation throughout. Representative images of Iba1 and IL-6 are shown in Figures 5 and 6, respectively.

**Figure 5.**
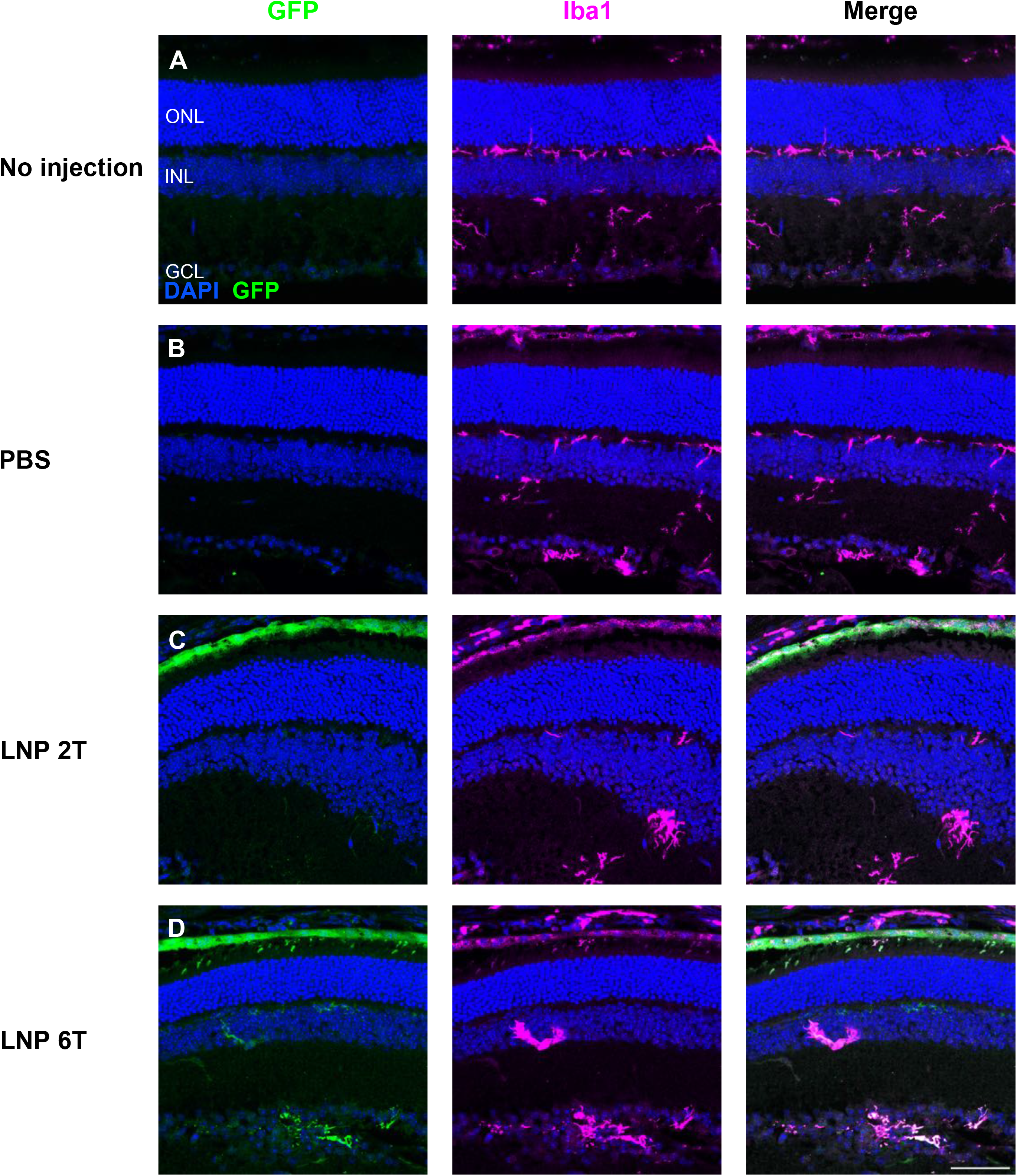
Immune surveillance by microglia is unchanged after intravitreal injections with EGFP mRNA LNPs. Microglia distribution, labelled with Iba1, was compared for 4 different conditions: (A) no injection, (B) PBS intravitreal injection, (C) EGFP mRNA 2TLNP intravitreal injection, or (D) EGFP mRNA 6T LNP intravitreal injection. Microglia are present in similar patterns in the outer plexiform layer (OPL), inner plexiform layer (IPL), and GCL for all conditions 6 hours post-injection. Images shown are representative of at least 3 retinas and are MIPs (scale bar: 50 μM).

**Figure 6.**
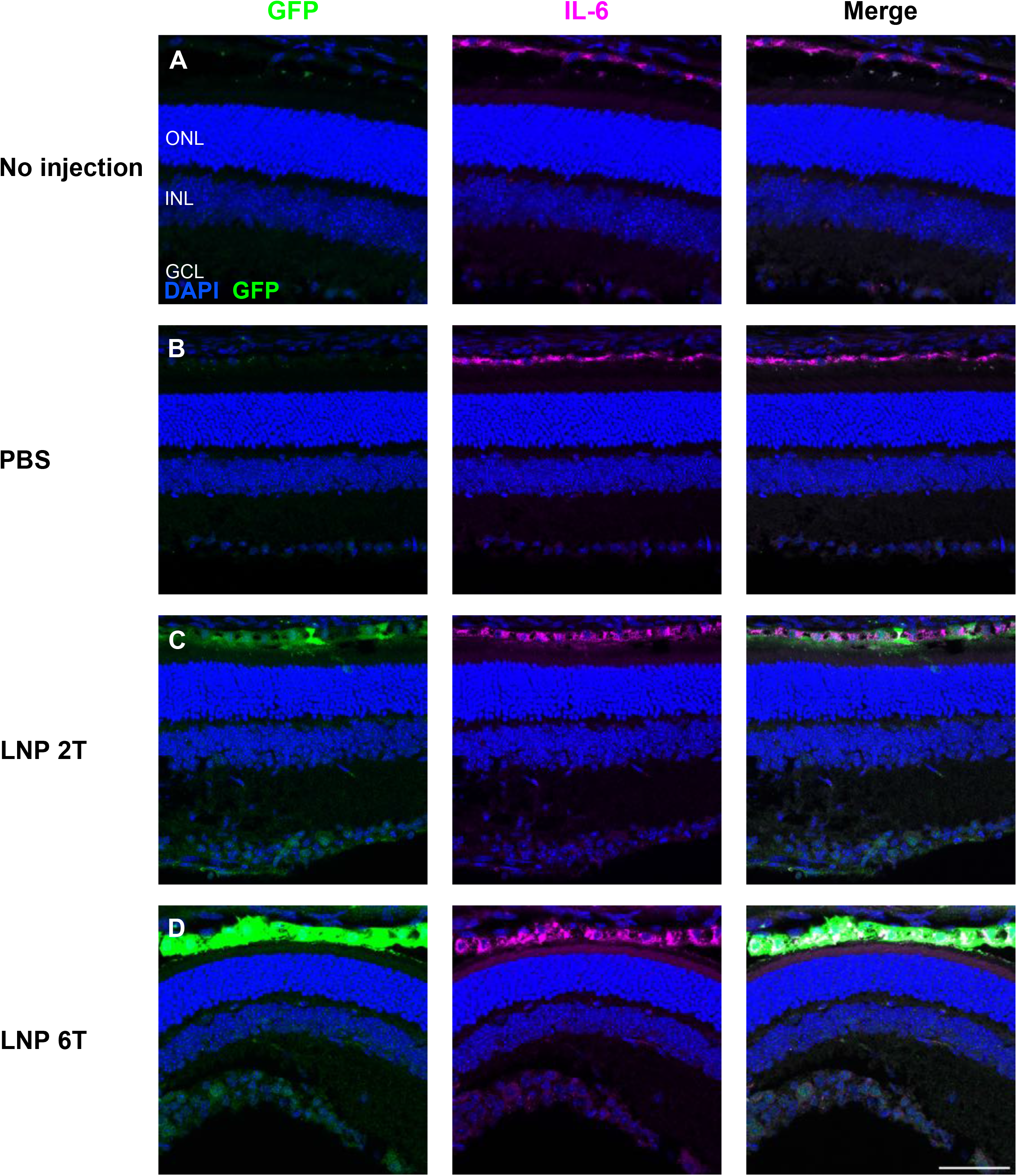
Intravitreal injections of EGFP mRNA LNPs does not affect the localization of cytokine IL-6. Expression of interleukin-6 in the RPE was similar comparing 4 conditions: (A) no injection, (B) PBS controls, (C) EGFP 2TLNP, or (D) EGFP 6T LNP at 6 hours post-intravitreal injection in mouse retinas. Images shown are representative of at least 3 retinas and are MIPs (scale bar: 50 μM).

### Potential Therapeutic Targets to Treat Retinal Disorders Using mRNA LNPs

Based on these preliminary results of the ability to deliver mRNA and increase protein expression in Müller glia and RPE cells specifically, there is the opportunity to further investigate therapeutic agents for delivery using mRNA LNP technology in future studies. A literature review of proteins that are deficient in Müller glia or RPE in inherited visual disorders found on RetNet are presented in Supplementary Tables 1 and 2. For example, replacement therapy of RLBP1 targeting Müller glia to treat recessive retinitis pigmentosa or TIMP3 in RPE cells to treat Sorsby’s fundus dystrophy could be explored further in future studies.

### Studies of mRNA LNP Delivery and Transfection in Human Fetal RPE and Adult Post-mortem Retina

Additional studies were conducted to determine transfection efficacy and cell-type specificity of mRNA LNPs in human retinas: human fetal RPE (fRPE) tissue and adult human retina and RPE. Human fRPE cells were dissected, cultured for 6 weeks, then treated with diluent only or 0.1 mg/mL, 0.25 mg/mL, or 0.5 mg/mL EGFP mRNA (2T) for 2 hours, then washed with fresh media and incubated for 16-24 hours before confocal imaging. RPE cells that were treated with mRNA LNPs had higher expression of EGFP than those treated with diluent only (Figure 7). The Corrected Total Cell Fluorescence (CTCF) is shown as a violin plot, and an ANOVA test with post-hoc Tukey multiple comparisons demonstrate a statistically significant difference between diluent only vs LNP treatment groups (Figure 7D). There is also nuclear expression of EGFP that overlaps with DAPI staining in the highest EGFP mRNA dose used (0.5 mg/mL). The pattern of cytoplasmic EGFP expression is similar in fRPE cells treated with 0.1 mg/mL and 0.25 mg/mL mRNA LNP (Supplementary Figure 3).

**Figure 7.**
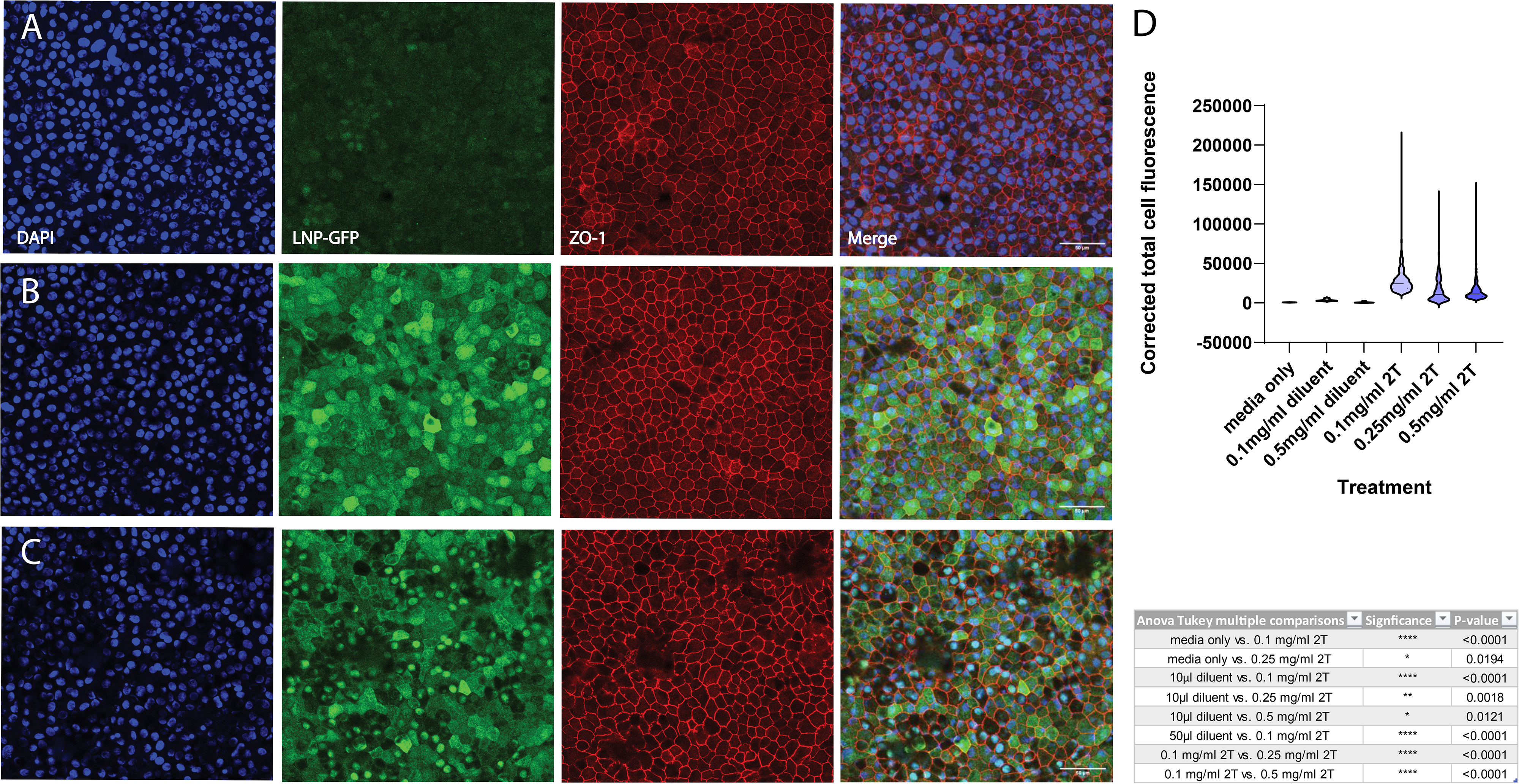
EGFP mRNA transfects human fRPE *in vitro*. Human fRPE cells treated with EGFP mRNA encapsulated in 2TLNPs for 2 hours express EGFP. (A) fRPE cells treated with diluent only (same volume as (B)) have minimal EGFP expression. (B) Cells were treated with 0.1 mg/mL mRNA LNP and have higher GFP expression than those treated with diluent only. (C) fRPE cells treated with 0.5 mg/mL mRNA LNP also have EGFP expression higher than (A) and additionally have nuclear EGFP expression. Images shown are representative of at least 3 cell culture wells and are MIPs (scale bar: 50 μM). (D) EGFP expression was quantified and plotted in a violin plot as Corrected Total Cell Fluorescence (CTCF). P-values from an ANOVA test with post-hoc Tukey multiple comparisons are shown in the table. CTCF values are significantly higher in LNP-treated cells than in diluent-treated cells.

Retinal explants from post-mortem adult globes were incubated with 0.1 mg/mL EGFP mRNA (2T or 6T LNP formulations) or diluent control for 4 hours, then placed in media before fixing, processing, and antibody staining for confocal imaging cross-sections and flat mounts. Human retinal cross-sections or flat mounts were stained with DAPI, anti-GFP, or anti-glial fibrillary acidic protein (GFAP). Retinal tissue incubated with EGFP mRNA in 6T LNPs express GFP in retinal cells that extend processes to the ONL and GCL (Figure 8A), a similar cellular morphology to the Müller glia that express GFP in the CD-1 mouse retina experiments. Cells incubated with EGFP mRNA 6T LNPs also expressed GFP in the fovea (Figure 8C) and in perivascular cells near blood vessels (Figure 8E), as indicated with arrows. Human retinal explants treated with EGFP mRNA LNPs 2Tyielded similar results: GFP was expressed in cells that extend apical and basal processes and surrounding blood vessels (Figure 8B) and near blood vessels in the retina near the RPE/choroid (Figure 8D).

**Figure 8.**
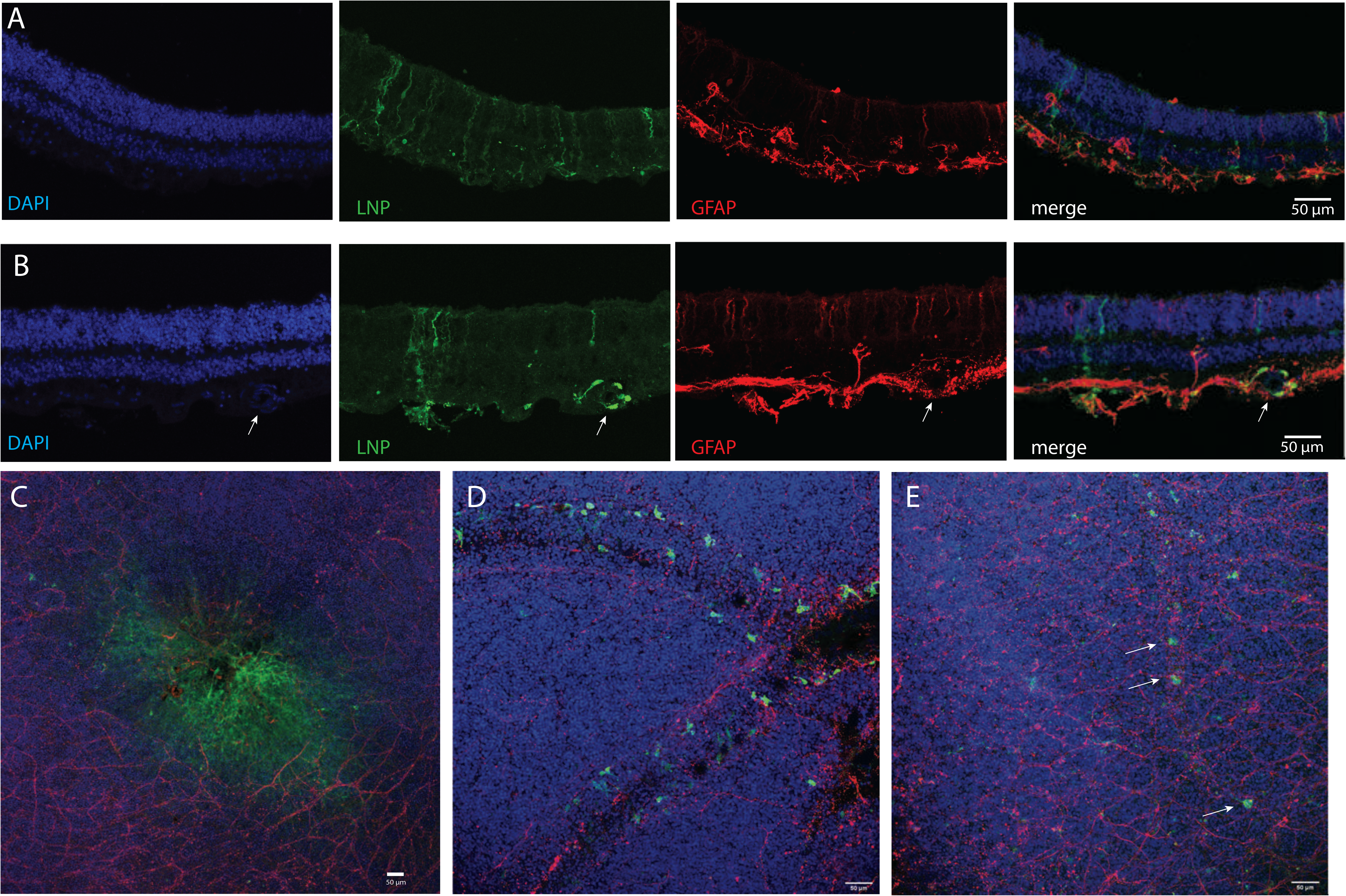
EGFP mRNA encapsulated in LNPs transfects adult human retina explants. Adult human post-mortem eye globes were treated with EGFP mRNA LNPs for 4 hours. (A) Confocal images of human retina cross-section treated with EGFP mRNA encapsulated in 6T LNPs and then stained with DAPI, anti-GFP, and anti-GFAP. GFP is expressed in cells with apical and basal processes. (B) Cross-section of human retina treated with EGFP mRNA encapsulated in 2T LNPs. GFP is expressed in perivascular cells surrounding a blood vessel, indicated by the arrow, and in cells with apical and basal processes. (C) Flat mount of parafoveal retina treated with EGFP mRNA in 6T LNPs. GFP is expressed in cells in the fovea. (D) Flat mount of retina with RPE and choroid treated with EGFP mRNA in 2T. Perivascular cells express GFP. (E) Flat mount of retina treated with 6T. Arrows indicate GFP positive cells along a small blood vessel. Images shown are representative of retinas and RPE from two globes from a single donor (scale bars: 50 μM).

## Discussion

Gene therapies are a promising treatment for monogenic retinal disorders. IRDs can lead to retinal degeneration and eventual vision impairment or loss, which has a significant impact on quality of life. Due to the lack of the retina’s ability to regenerate, earlier intervention and treatment is important to prevent irreversible damage. Currently, most potential treatments being investigated utilize AAVs or lentiviral vectors to deliver gene amplification. The only FDA-approved gene therapy for a visual disorder is voretigene neparvovec, an AAV2-based RPE65 replacement therapy for LCA^9^. While extremely promising, viral vectors have limited packaging capacity and have raised concerns about immunogenicity and genotoxicity, with limited ability to deliver larger transgenes or perform repeated injections^23^. In this study, we tested a non-viral method of gene augmentation therapy in retinas: mRNA encapsulated in LNPs. We successfully delivered EGFP mRNA LNPs to mouse retinas *in vivo* using intravitreal injections and demonstrated the ability to transfect primarily RPE and Müller glia cells. The kinetics of EGFP expression was rapid, seen as early as 6 hours and lasting at least 2 weeks. EGFP was expressed in retinal cells with similar tropism and kinetics when using LNPs that were either ApoE-and LDLR-dependent (2T) or LDLR-independent (6T). Furthermore, mRNA LNPs were able to transfect mouse Müller glia cells with a potential therapeutic agent, CNTF, demonstrating the ability to deliver a protein besides EGFP.

LNP-encapsulated mRNA delivery has been studied in various organ systems, including in the retina, liver, kidney, cerebral cortex, and red blood cells^14,15,24–29^. Clinically, the first use of mRNA LNP technology were two mRNA SARS-CoV-2 vaccinations developed during the COVID-19 pandemic^30,31^. In the retina specifically, previous studies examined the delivery of mCherry and luciferase mRNA LNP to the mouse retina, with expression in RPE and Müller glia cells over 4- 96 hours post-subretinal injection^14,15^. Our studies showed transfection in similar cell types in the mouse: RPE, Müller glia, and trabecular meshwork. Notably, we demonstrated the ability to transfect retinal cells for a longer duration, at least 2 weeks, and additionally were able to transfect the RPE using intravitreal injections, as opposed to the more invasive subretinal injections. This is particularly important in the context of mRNA LNPs; when compared to AAVs, the duration of transfection is shorter, thus possibly necessitating repeated administrations of treatment. Our ability to target the RPE with intravitreal injections is valuable; subretinal injections are often preferred for targeting the subretinal space but come with significant risks and would not be preferred for repeated injections: retinal detachment, vitreous hemorrhage, and choroidal neovascularization^32^. There are numerous RPE-specific deficiencies such as TIMP3 in Sorsby’s fundus dystrophy or BEST1 that causes macular dystrophy that could benefit from gene replacement therapy targeting the RPE, such as the mRNA LNPs in this study.

Our studies additionally studied the activation of the immune system to the mRNA LNP injections. We chose two immune markers to examine: IL-6, a cytokine that is upregulated and plays a key role during acute inflammation, and Iba1, a marker of microglia, immune surveillance cells in the retina^33,34^. Our results at a 6-hour timepoint showed no difference in IL-6 levels or Iba1 distribution between control groups no injection and PBS only intravitreal injection compared with EGFP 6T or 2T LNP intravitreal injections. These preliminary immune studies suggest that there is no significant immune system activation or reaction to the mRNA LNPs in the mouse eyes 6 hours post-treatment. Further studies including more cytokines and components of the complement system could be done to examine acute and chronic innate and adaptive system involvement in response to mRNA LNP therapy.

To our knowledge, our studies are the first to effectively transfect human retina and RPE *ex vivo* using LNP-encapsulated mRNA. The ability to transfect human retinal and RPE cells expands our potential applications, both clinically and experimentally. LNPs could be used for targeted gene replacement therapy in the human RPE or retina. The safety profile of mRNA LNPs is more ideal than viral vectors since they are less immunogenic, are cleared from the body, and are not able to incorporate into the genome^17,35^. There is overlap between the cell-type specificity of mRNA LNPs in the mouse and human retina: in both systems, RPE are transfected and cells that have processes that extend apically and basally express GFP, characteristic of Müller glia. Perivascular cells in the retina, possibly macrophages, express GFP in both adult post-mortem globes and neonatal mouse retinas. Importantly, we also showed the ability to transfect cells in the human fovea, which is a structure absent in the mouse retina. Overall, these results in human retinal and RPE tissue demonstrate that LNPs can be used to reliably deliver mRNA to both mouse and human retinas.

Additionally, mRNA LNPs are an intriguing option for experimental applications, like for transient transfection of various cell lines. Human RPE and induced pluripotent-or embryonic stem cell-derived RPE are difficult to transfect using traditional plasmid DNA methods, such as FuGENE, at least partially due to low cell viability during these protocols^36–40^. The mRNA LNPs used in these studies transfected cells with over 50% efficiency and caused minimal cell death. Therefore, this is a promising alternative for reliable and effective transfection of RPE and possibly other difficult to transfect cell lines.

LNP-encapsulated mRNA technology is exciting for a variety of future experimental and therapeutic applications. The ability to modify both mRNA and LNP to have less immunogenic effects, combined with its inability to incorporate into the genome and cause mutations, offers a safer alternative to viral vector-mediated gene therapies. Repeated intravitreal injections can be considered if longer durations of expression are needed for specific clinical applications. The ability to target the RPE with intravitreal injections makes this a unique opportunity to use gene therapy in the subretinal space without the need for subretinal injections. Additionally, the Müller glia are involved in retinal regeneration pathways, and therefore, mRNA LNPs in this study that target these cells could be used to deliver factors that could regenerate the retina^41,42^. LNP formulations can also be adjusted to target different cell types in the retina or to potentially increase gene expression duration. Exciting experimental applications can be pursued as well, such as delivering Cas9 for CRISPR/Cas9 gene editing or Cre recombinase using mRNA LNPs *in vitro* or *in vivo*. These future studies will further our understanding of the effectiveness and safety of LNP-encapsulated mRNA in the eye and advance the field of retinal gene therapy.

## Acknowledgments

The authors thank LuLu Callies and Stella Xu for their assistance with intravitreal injections and members of the Cherry Lab for their feedback during manuscript preparation. Experiments were designed, executed, and analyzed by CZC, GLS, ALE, and TJC with input from AF and PGVM on LNP and mRNA design and IAG and BDRL on human tissue experiments. IAG, BDRL (University of Washington, Seattle, WA), and Lions VisionGift (Portland, OR) collected human samples. CZC, ALE, and TJC wrote the manuscript with revisions and feedback from all authors.

---

**Supplementary Figure 1.**
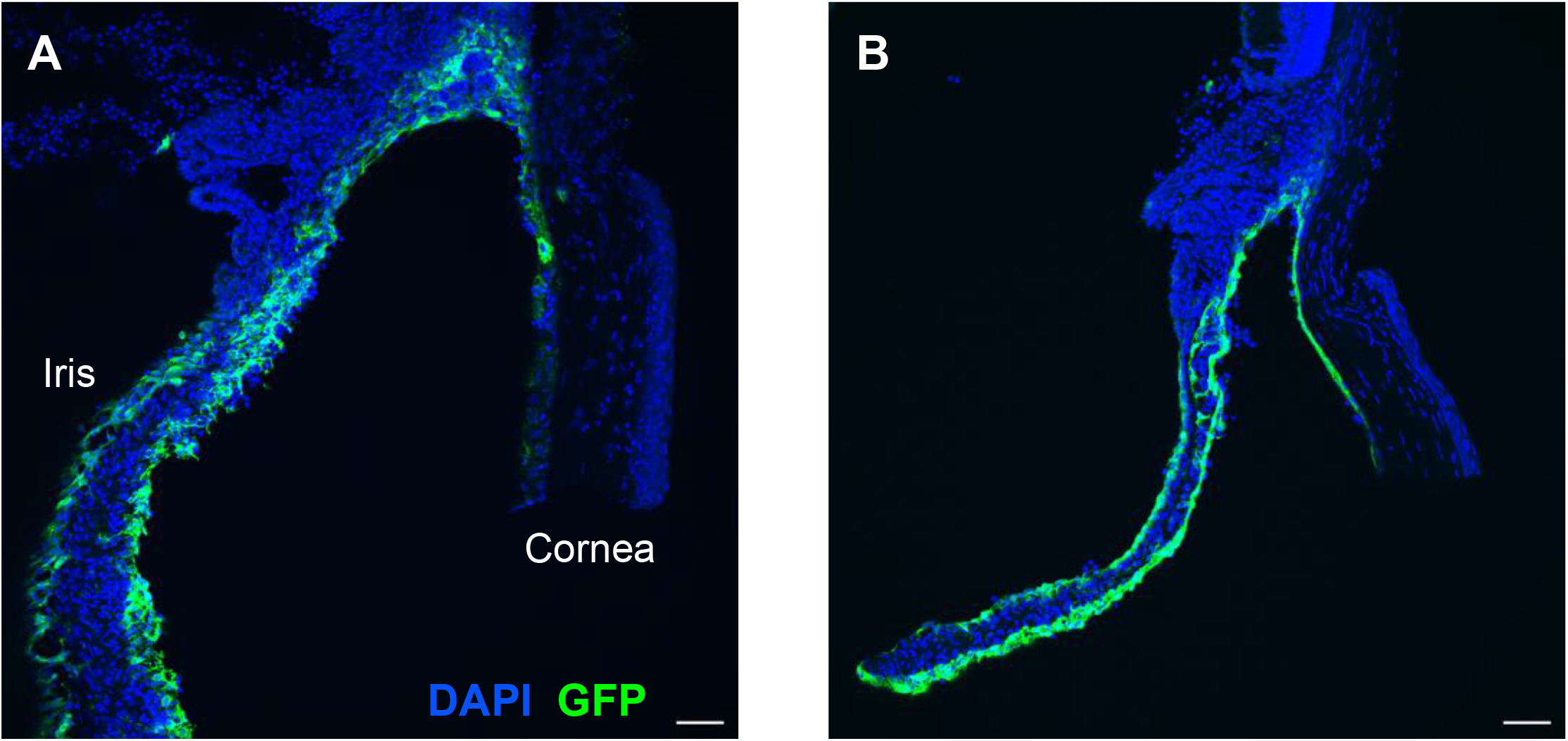
EGFP mRNA LNPs transfect the cornea and iris. (A) EGFP is expressed outside the retina in the cornea, iris, and trabecular meshwork in the iridocorneal angle 48 hours after intravitreal injection with EGFP mRNA encapsulated in 6T LNP. (B) Similarly, EGFP is expressed in the cornea, iris, and trabecular meshwork under the same experimental conditions as (A). Images shown are representative of at least 3 retinas and are MIPs (scale bar: 50 μM).

**Supplementary Figure 2.**
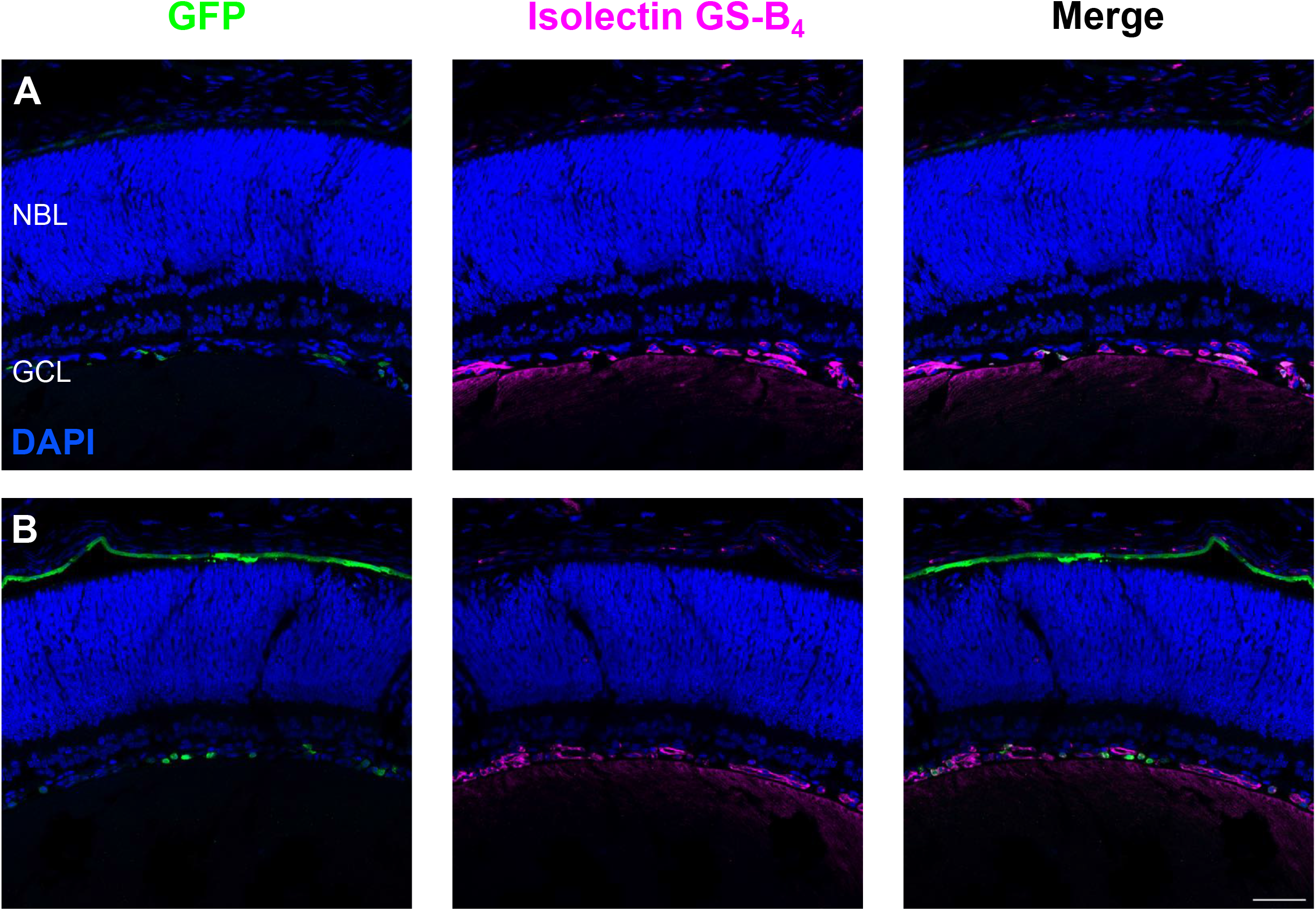
EGFP is expressed in perivascular and RPE cells in the neonatal mouse retina. Intravitreal injections of mouse pups (P0) were performed to deliver EGFP mRNA 2T LNPs. Retinas were harvested after 24 hours. (A) EGFP is co-expressed in the GCL near endothelial cells, labelled with Isolectin GS-B_4_. (B) EGFP is additionally expressed in the RPE region of the developing retina. Images shown are representative of at least 3 retinas and are MIPs (scale bar: 50 μM).

**Supplementary Figure 3.**
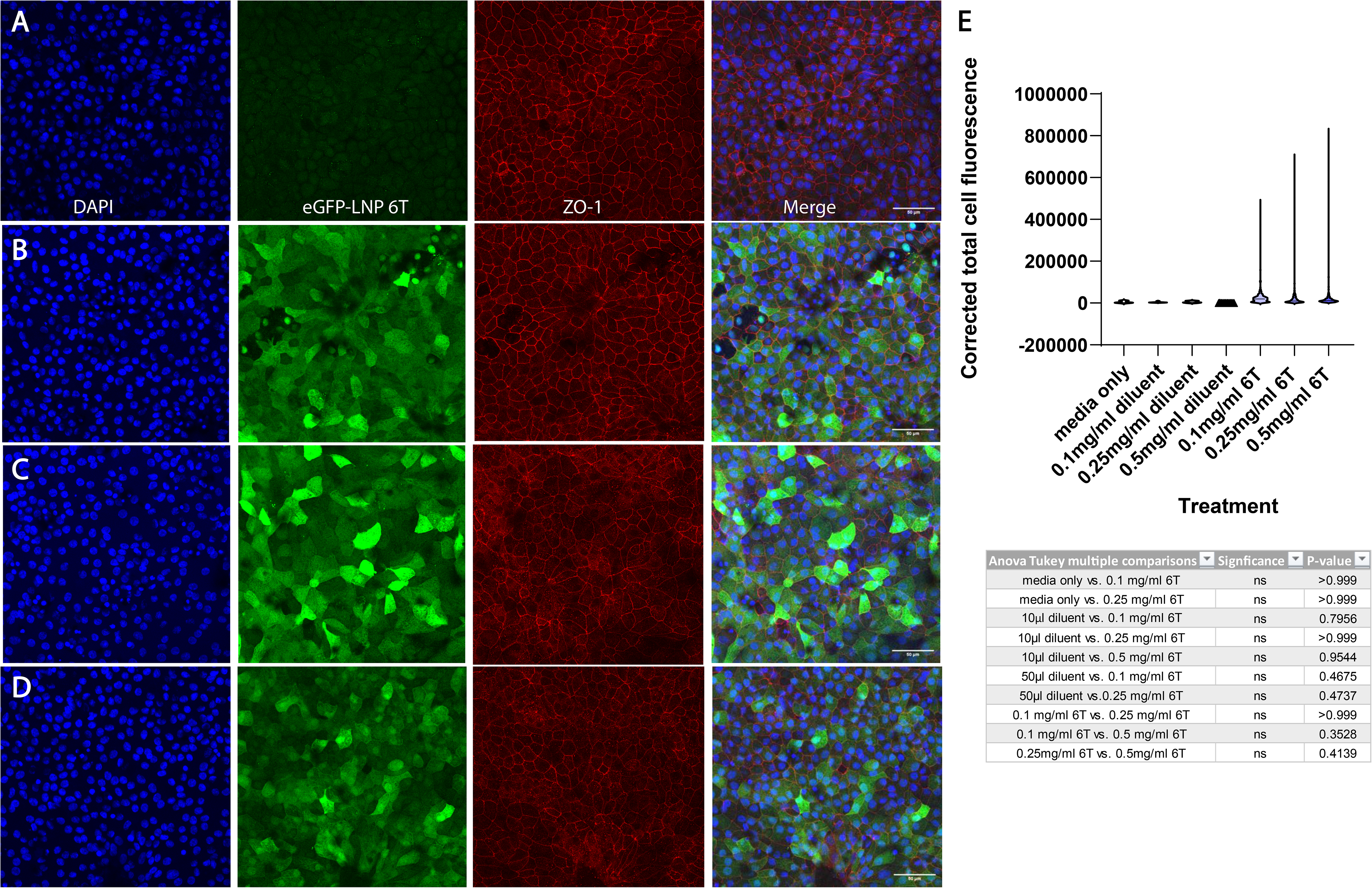
EGFP is expressed in fetal human RPE after treatment with 0.25 mg/mL EGFP 2T. (A) fRPE cells treated with diluent only (same volume as (B)) have minimal EGFP expression. (B) Cells were treated with 0.25 mg/mL mRNA LNP and have higher cytoplasmic GFP expression than those treated with diluent only. Images shown are representative of at least 3 cell culture wells and are MIPs (scale bar: 50 μM).

**Supplementary Table 1. Summary of potential therapeutic targets for mRNA LNP gene-replacement therapy in Müller cells.** This table lists IRDs that have mutations in proteins that are highly expressed in Müller cells. Specifically, candidate targets are presented with their gene of interest and protein name, chromosome location, relative expression level of the protein of interest in Müller glia, disease relevance, coding sequence size, Ensembl genome browser link, and link to a paper linking the mutation of interest to a retinal disorder.

**Supplementary Table 2. Summary of potential therapeutic targets for mRNA LNP gene-replacement therapy in RPE.** This table lists IRDs that have mutations in proteins that are highly expressed in RPE. Therapeutic targets are presented with their gene of interest and protein name, chromosome location, relative expression level of the protein of interest in Müller glia, disease relevance, coding sequence size, Ensembl genome browser link, and link to a paper linking the mutation of interest to a retinal disorder.

## Supplementary Material

Supplementary Figure 1

Supplementary Figure 2

Supplementary Figure 3

Supplementary Table 1

Supplementary Table 2

